# CREB regulates the expression of Type 1 Inositol 1,4,5-trisphosphate receptors

**DOI:** 10.1101/2021.05.05.442804

**Authors:** Vikas Arige, Lara E. Terry, Taylor R. Knebel, Larry E. Wagner, David I. Yule

## Abstract

Inositol 1,4,5-trisphosphate (IP_3_) receptors (IP_3_Rs) play a central role in regulating intracellular calcium signals in response to a variety of internal/external cues. Dysregulation of IP_3_R signaling is the underlying cause for numerous pathological conditions. It is also well established that the activity of IP_3_Rs is governed by several post-translational modifications including phosphorylation by protein kinase A (PKA). However, the long-term effects of PKA activation on expression of IP_3_R sub-types, remains largely unexplored. In this report, we investigate the effect of chronic activation of PKA on expression of IP_3_R sub-types. We demonstrate that expression of IP_3_R1 is augmented upon prolonged activation of PKA or upon ectopic over-expression of CREB but does not alter IP_3_R2 and IP_3_R3 sub-type abundance. Conversely, inhibition of PKA or blocking endogenous CREB diminished IP_3_R1 expression. We also demonstrate that agonist-induced Ca^2+^-release mediated by IP_3_R1 is significantly attenuated upon blocking endogenous CREB. Moreover, CREB by regulating the expression of KRAS-induced actin-interacting protein (KRAP) ensures proper localization and licensing of IP_3_R1. Overall, we report a crucial role for CREB in governing both the expression and proper localization of IP_3_R1.

**Summary statement:** We report a critical role of CREB in regulating the expression and proper localization of IP_3_R1. Agonist-induced Ca^2+^ release and Ca^2+^ puffs generated by IP_3_R1 are diminished upon blocking the function of endogenous CREB.

## INTRODUCTION

Inositol 1,4,5-trisphosphate (IP_3_) receptor (IP_3_R)-mediated redistribution of Ca^2+^ from the endoplasmic reticulum to the cytosol and other organelles occurs upon binding of IP_3_ (and the co-agonist Ca^2+^). IP_3_ is formed in response to a variety of hormones, growth factors, or neurotransmitters that activate phospholipase C isoforms (Berridge et al., 2000; Bootman et al., 2002; Clapham, 2007). The subsequent increase in intracellular Ca^2+^ regulates a myriad of physiological processes including secretion, contraction, gene transcription, proliferation, apoptosis, oxidative phosphorylation, learning and memory (Berridge et al., 2000; Clapham, 2007; Decuypere et al., 2011; Foskett et al., 2007). The human genome harbors three genes for IP_3_Rs: *ITPR1, ITPR2*, and *ITPR3* encoding IP_3_R1, IP_3_R2, and IP_3_R3 isoforms, respectively (Berridge, 1993; Bezprozvanny, 2005; Blondel et al., 1993; Furuichi et al., 1989; Sudhof et al., 1991). Each isoform is approximately 300 kDa and share 60-70% sequence similarity and are assembled as homo and hetero-tetrameric Ca^2+^ channels which are predominantly localized to the endoplasmic reticulum (Mignery et al., 1989; Monkawa et al., 1995; Wojcikiewicz and He, 1995). IP_3_R sub-types differ in their binding affinities for IP_3_ (Iwai et al., 2007; Newton et al., 1994) and the activity of individual subtype is differentially modulated by various post-translational modifications including phosphorylation by PKA (Masuda et al., 2010), PKC, CaMKII (Ferris et al., 1991b), AKT (Khan et al., 2006), ubiquitination (Alzayady and Wojcikiewicz, 2005), glycosylation (Rengifo et al., 2007), redox state of cells (Joseph et al., 2018), binding to ATP (Bezprozvanny and Ehrlich, 1993), and interaction with other proteins (Prole and Taylor, 2016).

IP_3_Rs are ubiquitously expressed; nevertheless, their expression is regulated in a tissue- and development-specific manner (De Smedt et al., 1997; Newton et al., 1994). For example, IP_3_R1 is predominantly expressed in the central nervous system while IP_3_R2 and IP_3_R3 are enriched in the peripheral tissues (Alzayady et al., 2013; Furuichi et al., 1989; Wojcikiewicz, 1995). IP_3_R1 knockout mice predominately die *in utero* or very early during post-natal development as a result of severe ataxia and seizures (Matsumoto et al., 1996; van de Leemput et al., 2007). Individual IP_3_R2 or IP_3_R3 knockout mice develop normally with subtle phenotypes; however the compound IP_3_R2/IP_3_R3 knockout animal displays severe defects in salivary, pancreatic and lacrimal exocrine secretion and die shortly after weaning due to their inability to assimilate macronutrients (Futatsugi et al., 2005). Moreover, perturbations in IP_3_R signaling or mutations in IP_3_Rs (Foskett, 2010; Terry et al., 2020) are associated with several pathological conditions including Alzheimer’s disease (Cheung et al., 2010), Huntington’s disease (Tang et al., 2005), schizophrenia (Tsuboi et al., 2015), amyotrophic lateral sclerosis (van Es et al., 2007), spinocerebellar ataxia (Ando et al., 2018; Ogura et al., 2001), Gillespie syndrome (Gerber et al., 2016), anhidrosis (Klar et al., 2014), diabetes (Roach et al., 2006) and cancer (Ueasilamongkol et al., 2020).

Previous reports also indicate that the number of IP_3_Rs can be acutely regulated. For example, Ca^2+^ influx through the L-type Ca^2+^-channels or NMDA receptors augment the expression of IP_3_R1 in neuronal cultures (Genazzani et al., 1999; Graef et al., 1999). Similarly, AP-1 (cFos/cJun dimer) and NFATc4 augmented IP_3_R1 expression in mouse cerebral cortical neurons (Mizuno et al., 2012; Mizuno et al., 2015). Box-I (corresponding to -334 to -318 bp region) in the *cis*-promoter region of IP_3_R1 was shown to be required for IP_3_R expression in N2a cells (Konishi et al., 1997). Moreover, 12-O-tetradecanoylphorbol-13-acetate or serum and 1,25-dihydroxyvitamin D3 or 17β-estradiol were shown to activate and inhibit the IP_3_R1 promoter activity, respectively (Kirkwood et al., 1997). Likewise, TGF-β is known to increase the phosphorylation of IP_3_R1 while inhibiting its expression in mesangial cells (Sharma et al., 1997). TGF-β is also known to inhibit the expression of IP_3_R1and IP_3_R3 in pre-glomerular afferent arteriolar smooth muscle cells (Pacher et al., 2008). We previously reported that TNF-α augments the activity and expression of IP_3_R1 via the transcription factor Sp1 (Park et al., 2008; Park et al., 2009). Nuclear Factor, Erythroid 2-Like 2 (NRF2) by binding to musculo-aponeurotic fibrosarcoma recognition element (MARE) in the IP_3_R3 promoter inhibits the expression of IP_3_R3 in both rat and human cholangiocytes (Weerachayaphorn et al., 2015). At the post-transcriptional level, miR-133a and miR-506 inhibit the expression of IP_3_R2 and IP_3_R3, respectively (Ananthanarayanan et al., 2015; Drawnel et al., 2012). The expression of IP_3_Rs is also induced by methamphetamine and cocaine exposure (Kurokawa et al., 2012). Nevertheless, additional molecular factors regulating the expression of IP_3_Rs remain largely underexplored.

Although several reports have clearly established the short-term/acute effects of raising cAMP on both the activity and phosphorylation status of IP_3_Rs (Ferris et al., 1991a; Patel et al., 1999; Soulsby and Wojcikiewicz, 2005; Wagner et al., 2008; Wagner et al., 2004; Wagner et al., 2003) any long term consequences of Protein kinase A (PKA) have not been defined. In the present study, we examined the long-term effects of chronic activation of PKA on the expression of IP_3_Rs in HEK-293 cells. Our results indicate that the endogenous protein levels of IP_3_R1 are augmented in response to increasing PKA activity by exposure to forskolin, while the levels of IP_3_R2 and IP_3_R3 remain unaffected. We also show that this effect is mediated via the transcription factor CREB. Consistent with this observation, ectopic over-expression of constitutively active CREB in HEK-293 cells augmented the protein levels of IP_3_R1 and KRAP, a binding partner of IP_3_R important for localization and function (Fujimoto et al., 2011b). This transcriptional regulation of IP_3_R1 abundance had functional consequences as dominant negative CREB transfection significantly diminished agonist-induced Ca^2+^ release and Ca^2+^ puffs, the elementary calcium signals generated by IP_3_Rs. Taken together, our results indicate a crucial role for CREB in governing the expression of IP_3_R1 through its endogenous promotor and subsequently the underlying agonist-evoked Ca^2+^ release/Ca^2+^puffs by IP_3_R1 in HEK-293 cells.

## RESULTS

Previous results from our laboratory and other groups have conclusively documented that all the three IP_3_Rs are phosphorylated by PKA (Wagner et al., 2008; Wojcikiewicz and Luo, 1998). The sites of phosphorylation and the acute functional consequences of phosphorylation of IP_3_R activity have also been extensively studied (Soulsby et al., 2004). Nevertheless, the long-term effects of PKA activation on the endogenous protein levels of IP_3_Rs (stimulus-transcriptional coupling) have not been fully determined.

### Forskolin augments the expression of IP_3_R1 in HEK-293 cells

In order to investigate whether PKA activation has any long-term effects on the endogenous protein levels of IP_3_Rs, we treated HEK-293 cells with 10 μM forskolin for 12 hours to raise cAMP and activate PKA followed by Western blotting to determine the protein levels of IP_3_Rs. The endogenous protein level of IP_3_R1 significantly increased following treatment with 10 μM forskolin (Fig. 1A, B). In contrast, the endogenous protein levels of both IP_3_R2 (Fig. 1C, D) and IP_3_R3 (Fig. 1E, F) remained unaltered. Moreover, treatment with 10 μM forskolin augmented the endogenous IP_3_R1 protein levels in a time-dependent manner (Fig. S1). Consistently, the IP_3_R1 remained phosphorylated following 12 hours of treatment with 10 μM forskolin (Fig. S2). These results indicate that long-term treatment with forskolin augments both the endogenous IP_3_R1 protein levels and the extent to which the receptors remained phosphorylated in HEK-293 cells.

**Fig. 1:**
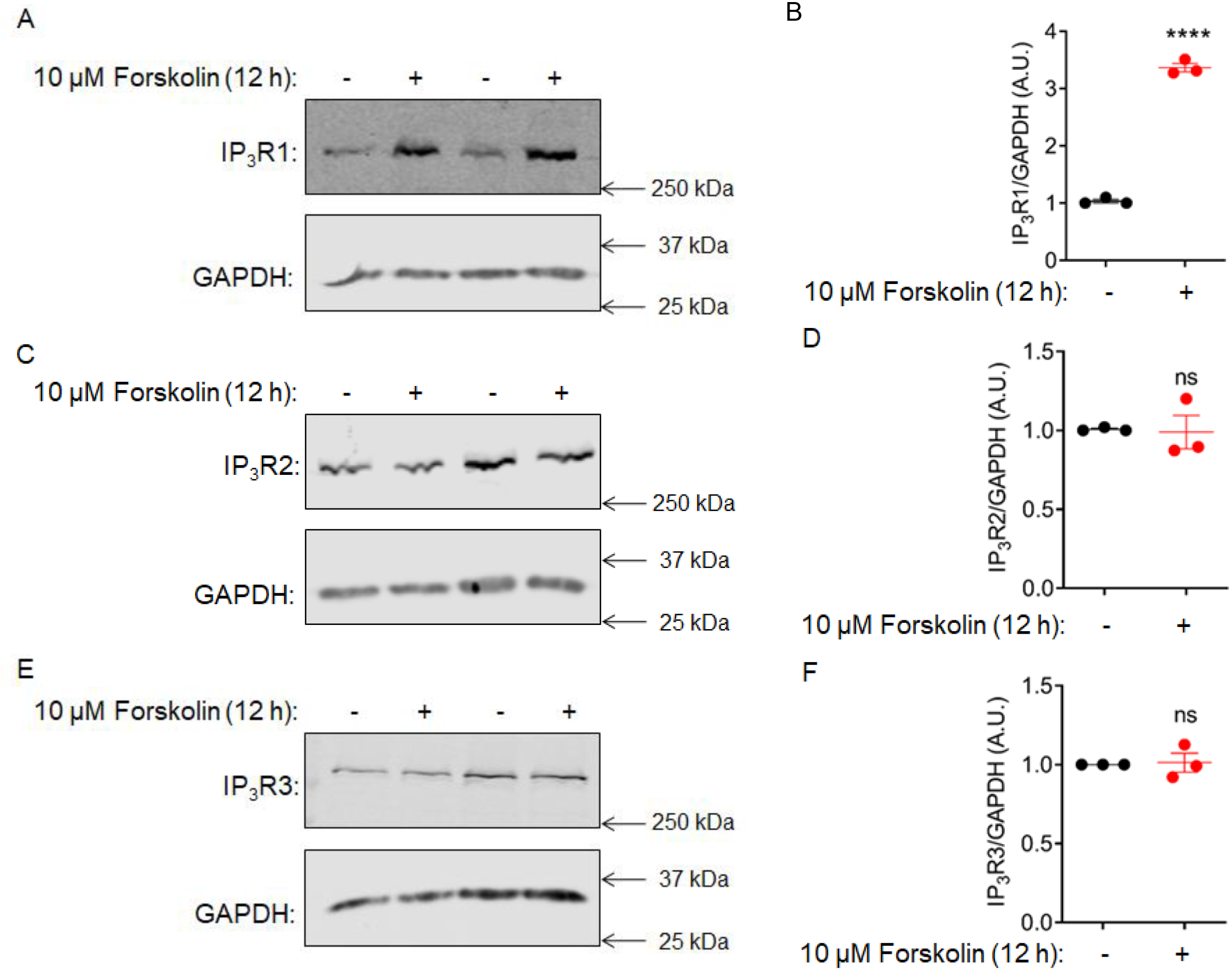
Forskolin augments the endogenous protein level of IP_3_R1 in HEK-293 cells. HEK-293 cells were treated with 10 μM forskolin for 12 hours followed by protein isolation. (A) A representative Western blot depicting forskolin-induced endogenous IP_3_R1 protein level in HEK-293 cells. (B) Quantification from three independent blots is shown. (C) Western blot depicting no effect of forskolin on the endogenous protein levels of IP_3_R2 in HEK-293 cells. (D) Quantification from three independent blots is shown. (E) Western blot depicting no effect of forskolin on the endogenous protein levels of IP_3_R3 in HEK-293 cells. (F) Quantification from three independent blots is shown. Statistical significance was determined by Student’s t-test (unpaired, two-tailed). ****p<0.0001. ns; not significant. (Lane numbers 1 & 2: 20 μg protein and lane numbers 3 & 4: 30 μg protein was loaded).

### *In silico* analysis revealed putative binding sites for CREB in the proximal promoter domains of IP_3_Rs

Forskolin activates adenylyl cyclase resulting in increased intracellular cAMP (Insel and Ostrom, 2003). Increased cAMP binds to the regulatory subunits of protein kinase A and one important consequence is that the free catalytic subunit translocates into the nucleus. In the nucleus, the catalytic subunit phosphorylates and activates the transcription factor CREB. Once phosphorylated, CREB recruits the transcriptional machinery required for transcription of target genes (Altarejos and Montminy, 2011). Since forskolin treatment augmented the endogenous protein level of IP_3_R1, we employed multiple *in silico* transcription factor prediction tools including Alibaba2.1, TFBIND, Consite, and PROMO which rely on different algorithms to predict transcription factor binding sites for CREB in the 1 kb proximal promoter domains of IP_3_Rs. Interestingly, three out of the four tools predicted at least one binding site for CREB in the proximal promoter domains of all the three IP_3_Rs (Fig. 2A). In addition to this, these tools predicted binding sites for transcription factors Sp1, Egr1, and NF-KB amongst others. This concurs with a previous report from our group wherein we demonstrated that Sp1 is involved in regulating the expression of IP_3_R1 (Park et al., 2009).

**Fig. 2:**
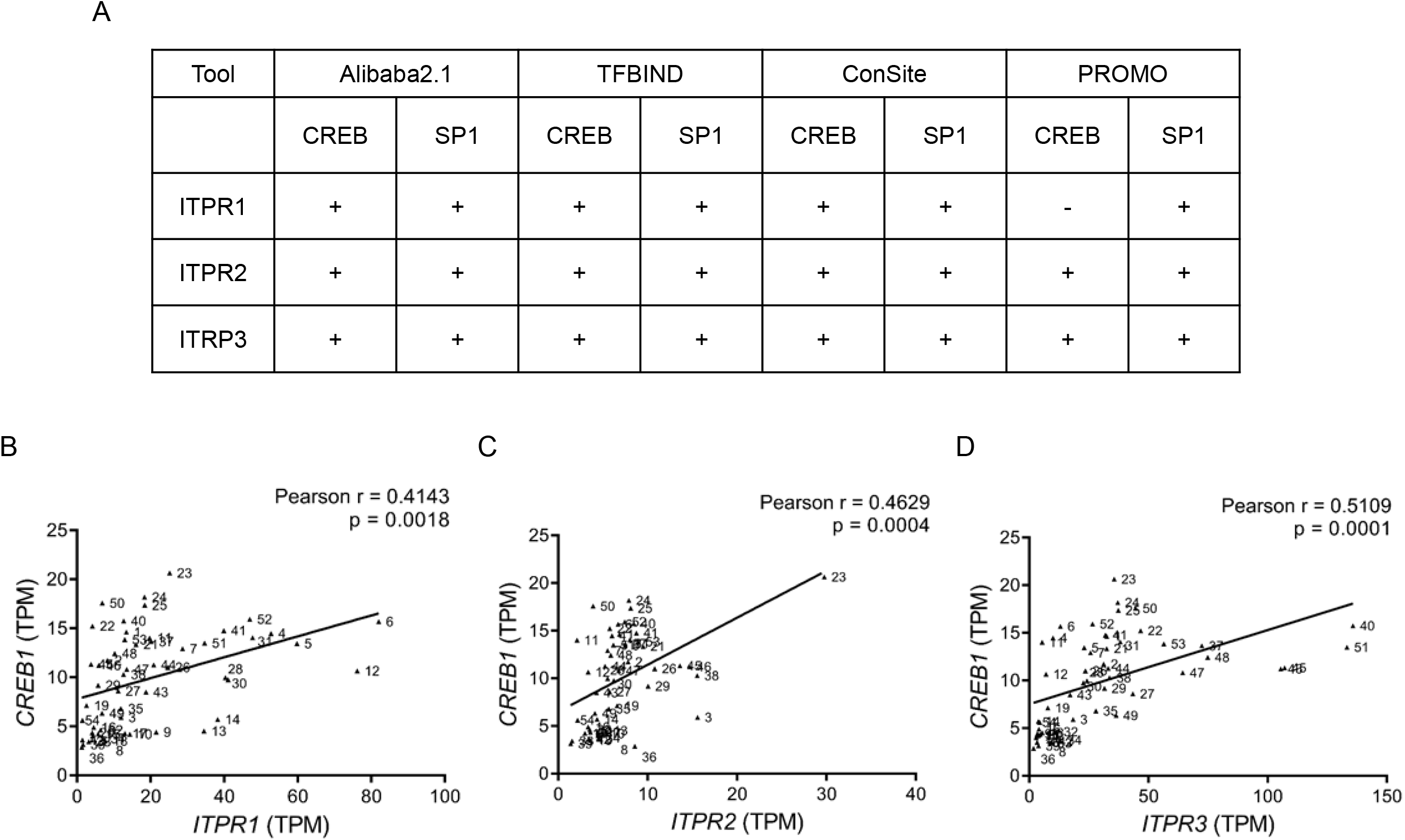
*In silico* analysis revealed putative binding sites for CREB in the proximal promoter domains of ITPR1/2/3 genes. (A) Table showing results from the computational analysis using various transcription factor prediction tools to predict putative binding sites for CREB and Sp1 in the 1 kb proximal-promoter domains of ITPR1/2/3 genes. (‘+’ and ‘-’ denote predicted and not predicted, respectively). RNAseq data mined from GTEx portal displayed significant positive correlation between the transcript levels of (B) *ITPR1* vs. *CREB*, (C) *ITPR2* vs. *CREB*, and (D) *ITPR3* vs. *CREB* across various human tissues (n=54). TPM; Transcripts per Million.

### Correlation between the transcript levels of IP_3_Rs and CREB across various human tissues

Next, to determine if there is any correlation between the transcript levels of IP_3_Rs and CREB, we mined transcriptomics data for CREB and IP_3_Rs from GTEx database, a publicly available database. The transcript levels of IP_3_R1, IP_3_R2, and IP_3_R3 displayed significant positive correlations with the transcript levels of CREB across various human tissues (n=54 tissue types) (Fig. 2B, C, D) (Table S1). Taken together with the *in silico* transcription factor predictions, these results indicate a possible role for CREB in regulating the expression of IP_3_Rs across various human tissues.

### Involvement of PKA-CREB axis in regulating the expression of IP_3_R1

To dissociate the acute effects of IP_3_R1 phosphorylation and to demonstrate the involvement of CREB in regulating the expression of IP_3_Rs, increasing amounts of the constitutively active VP16-CREB expression plasmid was transfected into HEK-293 cells followed by Western blotting to determine the endogenous protein levels of all the three subtypes of IP_3_Rs. VP16-CREB binds the cAMP response element (CRE) through the DNA binding domain of CREB and transcriptional activation is facilitated by the transcriptional activation domain of the herpes virus VP16 protein (Riccio et al., 1999). Interestingly, transient transfection of VP16-CREB to over-express CREB augmented the endogenous protein levels of IP_3_R1 (Fig. 3A, B) while the levels of IP_3_R2 and IP_3_R3 remained unaltered (Fig. S3A, B). In order to substantiate the role of CREB in regulating the expression of IP_3_R1, KCREB plasmid, a dominant negative form of CREB, was transfected into HEK-293 cells to block the function of endogenous CREB (Walton et al., 1992). In contrast to the over-expression of constitutively active CREB, Western blotting revealed KCREB transfection resulted in a concentration-dependent decrease in the endogenous IP_3_R1 protein levels in HEK-293 cells (Fig. 3C, D). These results indicate that IP_3_R1 is a *bona fide* target of CREB in HEK-293 cells.

**Fig. 3:**
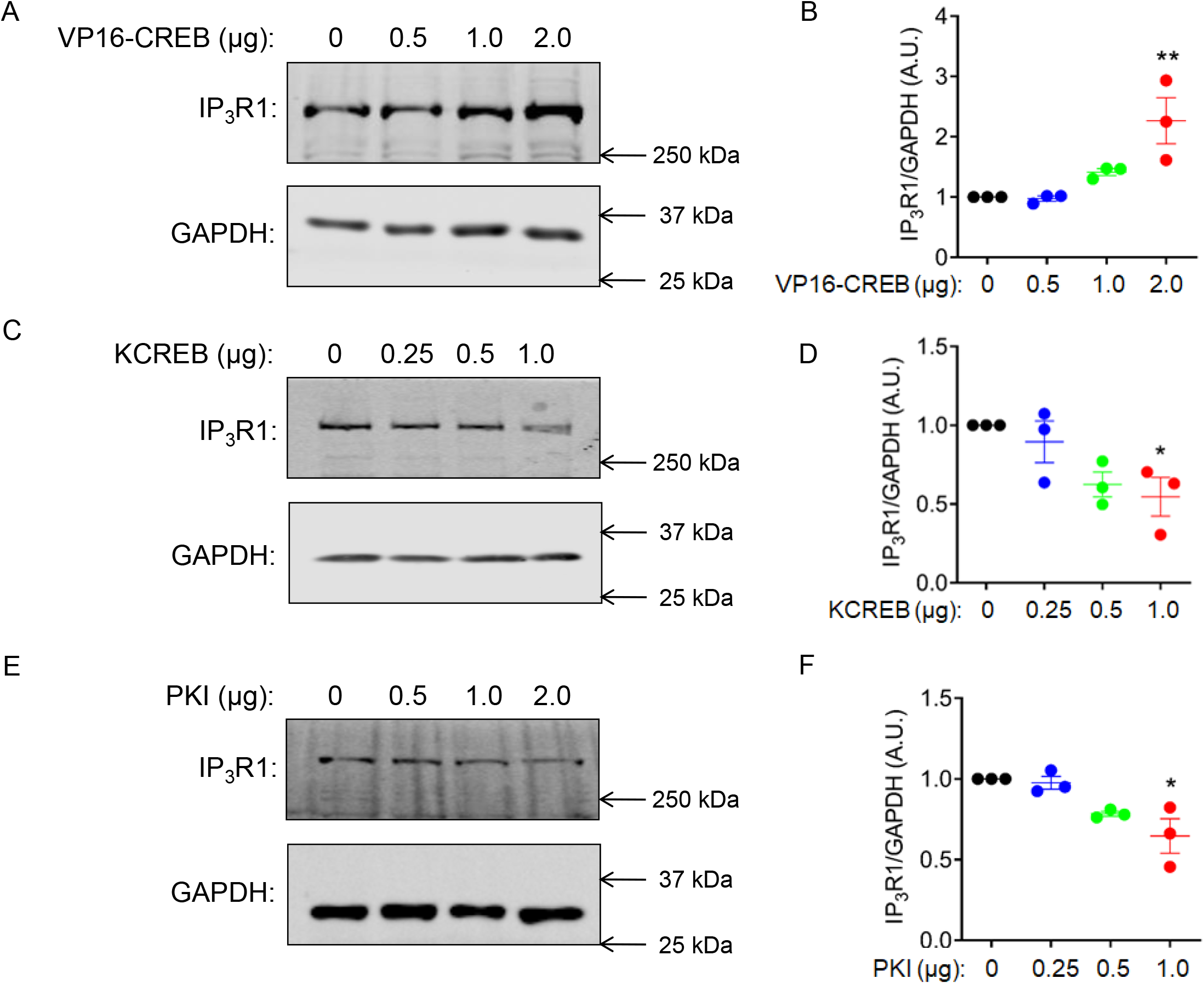
Evidence for involvement of PKA-CREB axis in governing the expression of IP_3_R1. HEK-293 cells were transiently transfected with increasing amounts of VP16-CREB expression plasmid and total protein was isolated 36 hours following transfection. (A) A representative blot showing over-expression of the VP16-CREB plasmid resulted in a dose-dependent increase in the endogenous IP_3_R1 protein levels in HEK-293 cells. (B) Quantification from three independent blots is shown. HEK-293 cells were transiently transfected with increasing amounts of KCREB plasmid and total protein was isolated 36 hours following transfection. (C) A representative Western blotting displaying the effect of blocking endogenous CREB, using the KCREB plasmid, on the endogenous IP_3_R1 protein levels in HEK-293 cells. (D) Quantification from three independent blots is shown. HEK-293 cells were transiently transfected with increasing amounts of PKI plasmid and total protein was isolated 36 hours following transfection. (E) A representative blot showing inhibition of endogenous PKA, using the PKI plasmid, resulted in diminished endogenous IP_3_R1 protein levels in HEK-293 cells. (F) Quantification from three independent blots is shown. Statistical significance was determined by one-way ANOVA with Tukey’s multiple comparisons post-test. *p<0.05, **p<0.01.

Since forskolin treatment activates PKA, to assess the role of PKA in regulating the expression of IP_3_R1 through CREB, a PKI plasmid (PKA inhibitor plasmid) (Day et al., 1989) was transiently transfected into HEK-293 cells followed by Western blotting to probe for endogenous IP_3_R1 levels. As expected, transfection of PKI plasmid resulted in a decrease in endogenous IP_3_R1 protein levels in HEK-293 cells (Fig. 3E, F). These results indicate involvement of PKA-CREB axis in regulating the expression of IP_3_R1 in HEK-293 cells.

### Forskolin-induced IP_3_R1 expression is mediated by CREB

In order to determine the involvement of CREB in mediating the effects of PKA activation, HEK-293 cells were transiently transfected with either pcDNA or KCREB followed by treatment with 10 μM forskolin and Western blotting. KCREB transfection resulted in decreased IP_3_R1 protein level while treatment with 10 μM forskolin resulted in enhanced IP_3_R1 protein level in HEK-293 cells (Fig. 4A, B). The forskolin-induced endogenous IP_3_R1 protein level was markedly diminished upon blocking endogenous CREB (Fig. 4A, B). These results further emphasize a role for CREB in forskolin-induced IP_3_R1 expression.

**Fig. 4:**
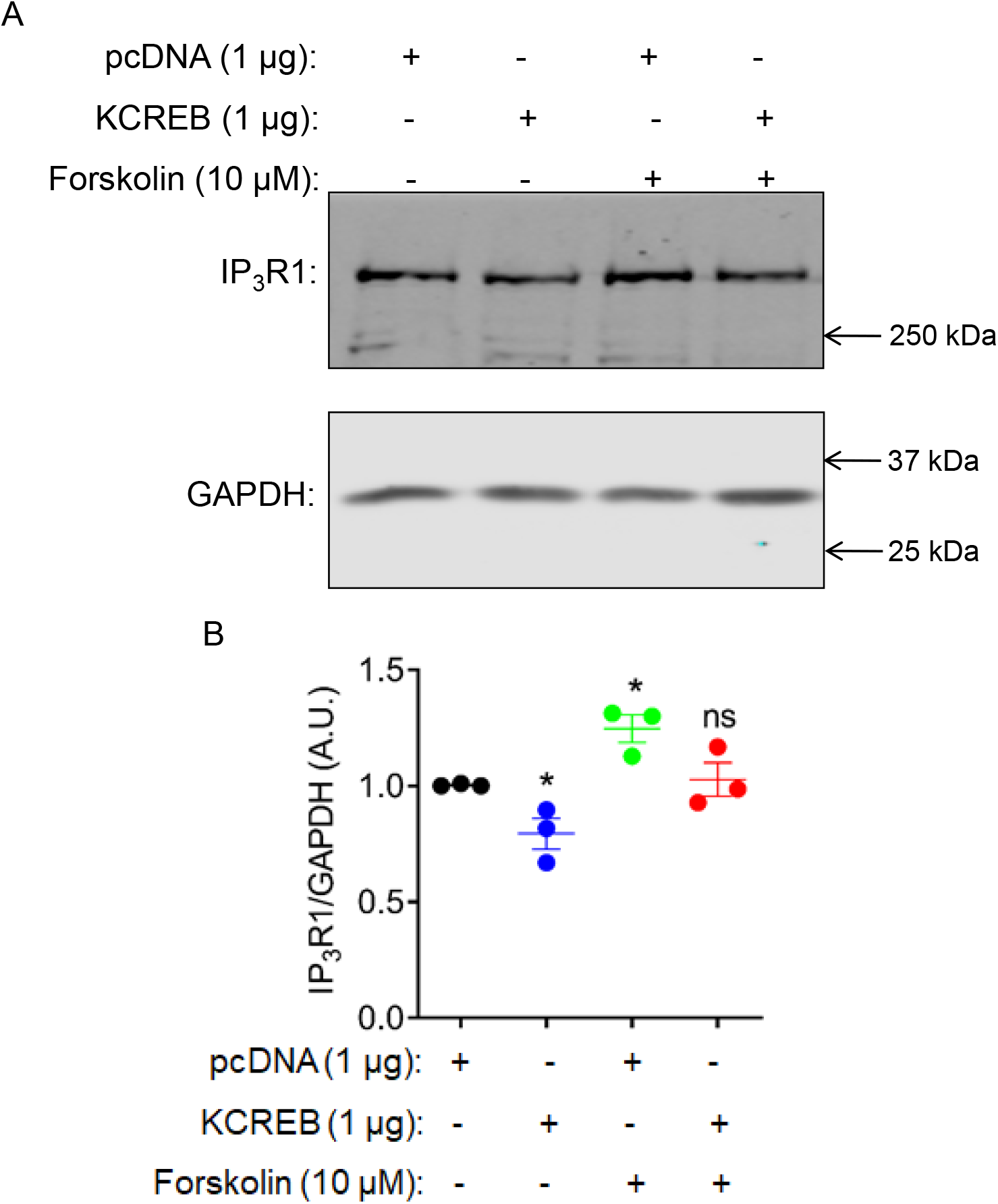
A role of CREB in augmenting forskolin-induced IP_3_R1 expression. HEK-293 cells were transiently transfected with either pcDNA or KCREB plasmid. After 24 hours transfection, the cells were treated with 10 μM forskolin (in the treatment group) or left untreated for additional 12 hours. Following 36 hours from transfection, protein was isolated for Western blotting. Western blotting performed using (A) HEK-293 cells treated with forskolin under basal conditions or upon blocking CREB, using the KCREB plasmid, suggests involvement of CREB in forskolin-induced IP_3_R1 expression. (B) Quantification from three independent blots is shown. Statistical significance was determined by Student’s t-test (unpaired, two-tailed). *p<0.05. ns; not significant.

### Agonist-induced Ca^2+^ release is markedly diminished upon blocking endogenous CREB in a population of cells

We next investigated the impact of CREB on agonist-induced Ca^2+^ release from HEK-293 cells engineered using CRISPR-Cas9 technology to only express IP_3_R1 at endogenous levels by the deletion of IP_3_R2/3 normally expressed in HEK-293 cells (Alzayady et al., 2016). These cells (designated hR1 endo cells) were utilized in all our subsequent experiments to prevent the contribution of IP_3_R2 and IP_3_R3 on Ca^2+^ release as a function of heterotetrameric populations of IP_3_Rs. These cells were transfected either with pcDNA or KCREB and Ca^2+^ release in response to the muscarinic agonist carbachol (CCh) was monitored using a Flexstation plate-reader assay platform with microfluidics. The increase in cytosolic Ca^2+^ in response to increasing concentrations of CCh (0.1-100 μM) was then determined. Notably, stimulation of a population of cells transfected with KCREB displayed a significant decrease in Ca^2+^ increase as compared to cells transfected with pcDNA at sub-maximal concentration of CCh (10 µm) (Fig. 5A). This data demonstrated that KCREB transfection in these cells resulted in decreased IP_3_R1 expression consistent with a decreased Ca^2+^ release in response to agonist stimulation.

**Fig. 5:**
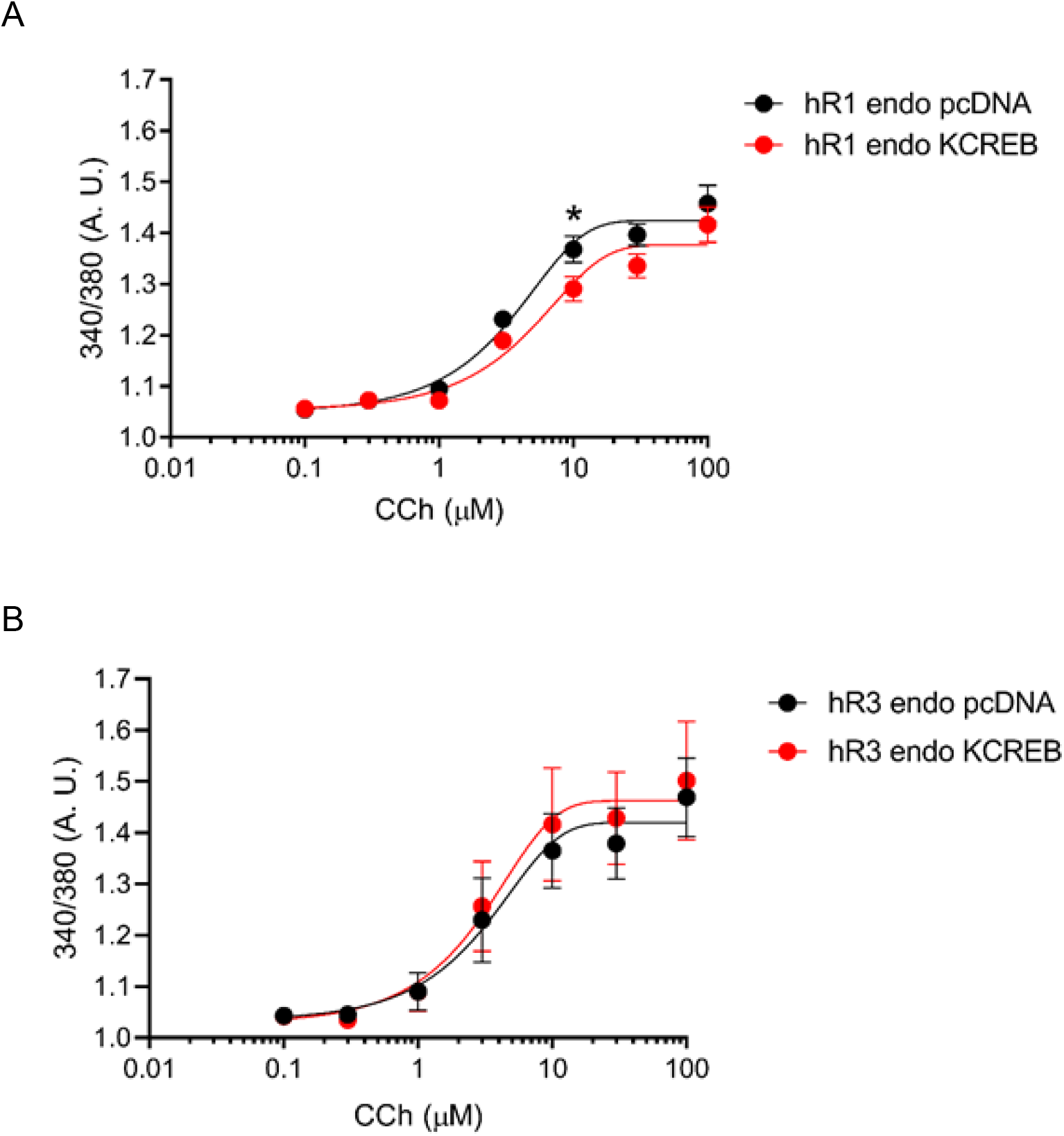
Inhibition of endogenous CREB diminished agonist-induced calcium release in a population-based calcium imaging assay. (A) HEK-293 IP_3_R2/3^-/-^ cells (hR1 endo cells) transiently transfected with either pcDNA or KCREB were loaded with Fura-2/AM and imaged using a Flexstation 96-well plate reader in response to increasing concentrations (0.1-100 μM) of CCh. The Ca^2+^ release at 10 μM CCh was significantly diminished in hR1 endo cells transfected with KCREB as compared to cells transfected with pcDNA. However, (B) HEK-293 IP_3_R1/2^-/-^ (hR3 endo cells) transfected with either pcDNA or KCREB subjected to a similar protocol, did not display such a difference in calcium release in response to increasing concentrations of CCh. Data presented as mean ± SEM. Statistical significance was determined by Student’s t-test (unpaired, two-tailed). *p<0.05.

In contrast, there was no significant difference in agonist-induced Ca^2+^ release from HEK-293 cells engineered using CRISPR-Cas9 technology to only express IP_3_R3 (Alzayady et al., 2016) (designated as hR3 endo cells) transfected with equal amounts of either pcDNA or KCREB (Fig. 5B). Indeed, these results are in agreement with our observations that the endogenous protein levels of IP_3_R2 and IP_3_R3 remain unaltered upon over-expressing CREB (Fig. S3A, B).

### CCh-induced Ca^2+^ release is markedly attenuated upon blocking endogenous CREB

Given that the agonist-induced Ca^2+^ release is inhibited in a population-based assay with presumably non-uniform transfection efficiency, we next performed single cell imaging using an epifluorescence microscope where transfected cells could be optically identified. For single cell imaging, hR1 endo cells were co-transfected with either pcDNA or KCREB and mCherry plasmids. Following 36 hours transfection, the cells were loaded with 2 μM Fura-2/AM and the changes in cytosolic Ca^2+^ in response to increasing doses of CCh were monitored. mCherry positive hR1 endo cells transfected with KCREB displayed much lower amplitudes (Fig. 6B, E) compared to mCherry positive pcDNA transfected cells (Fig. 6A, D) at various concentrations of CCh. Consistent with this, the average increase in amplitude and the peak amplitude changes were much lower in cells transfected with KCREB as compared to pcDNA (Fig. 6C, F, H). Moreover, the percentage of KCREB transfected cells responding to various concentrations of CCh at a specific threshold (F/F_o_>0.02) were diminished when compared to pcDNA transfected cells (Fig. 6G). Overall, these results provide mechanistic insights supporting a key role of CREB in governing the expression of IP_3_R1 and the extent of agonist-evoked Ca^2+^ release.

**Fig. 6:**
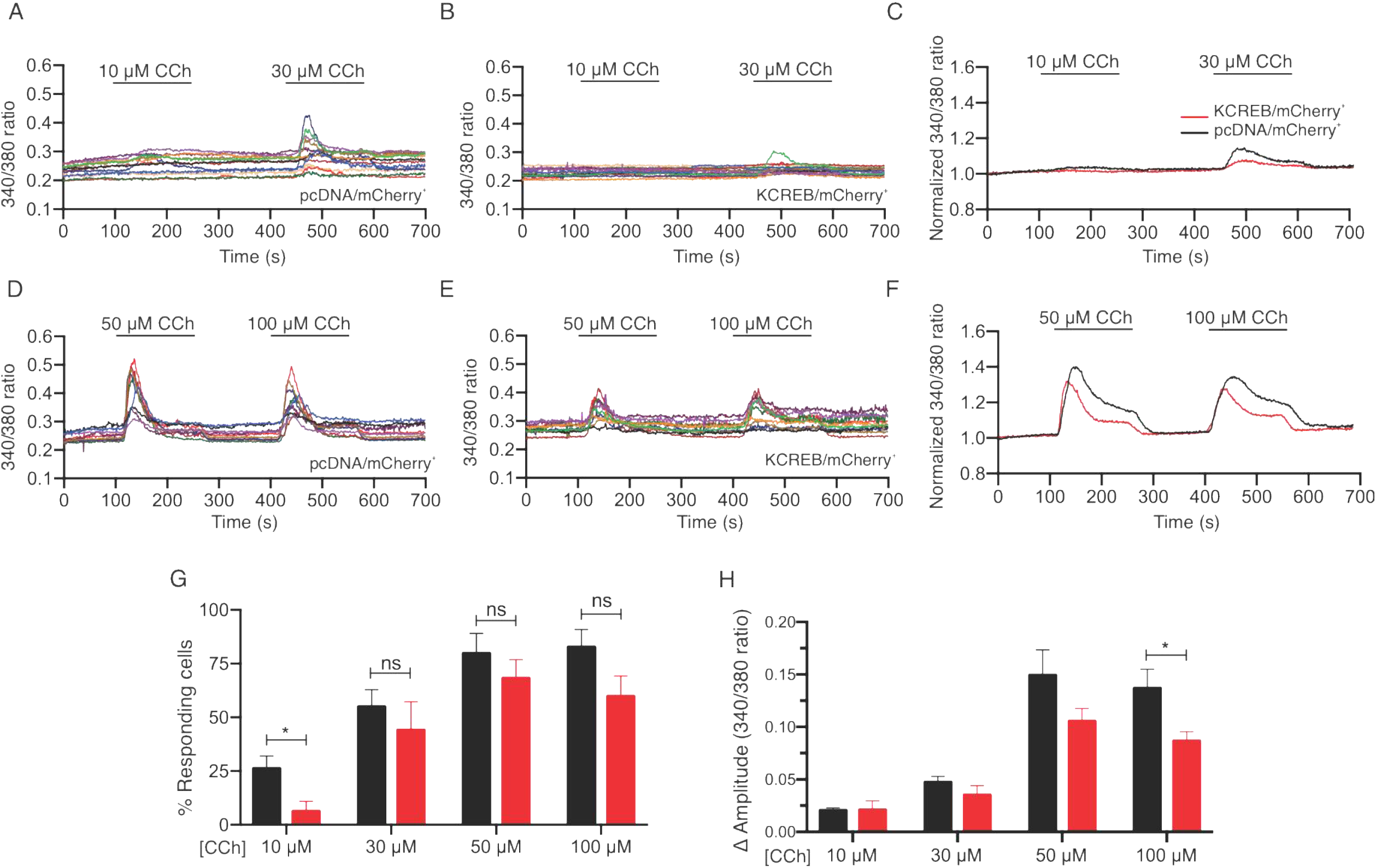
Inhibition of endogenous CREB diminished CCh-induced calcium release in single cells: hR1 endo cells transiently co-transfected with mCherry and either pcDNA or KCREB plasmids were loaded with Fura-2/AM at room temperature for 25 mins followed by imaging using an inverted epifluorescence microscope. Only those cells which were mCherry positive were considered for analysis. Representative traces showing increase in cytosolic Ca^2+^ from (A) hR1 endo cells transfected with pcDNA or (B) KCREB in response to CCh (10 and 30 μM). (C) A summary of average rise in cytosolic Ca^2+^ from hR1 endo cells transfected with pcDNA (n=48 cells) or KCREB (n=45 cells) in response to CCh (10 and 30 μM) is shown. Representative traces showing increase in cytosolic Ca^2+^ from (D) hR1 endo cells transfected with pcDNA or (E) KCREB in response to CCh (50 and 100 μM). (F) A summary of average rise in cytosolic Ca^2+^ from hR1 endo cells transfected with pcDNA (n=62 cells) or KCREB (n=50 cells) in response to CCh (50 and 100 μM) is shown. (G) Bar graph showing the percentage of responding cells (an amplitude change greater than 0.02, from 7 independent experiments) at various concentrations of CCh. Data presented as mean ± SEM. Statistical significance was determined by Student’s t-test (unpaired, two-tailed). *p<0.05. ns; not significant. (H) Bar graph showing change in amplitude (peak ratio – basal ratio: average of initial 20 ratio points)at various concentrations of CCh. Data presented as mean ± SEM. Statistical significance was determined by two-way ANOVA with Bonferroni test. *p<0.05. *p < 0.05.

### Diminished expression of IP_3_R1 results in dampened ci-IP_3_ evoked Ca^2+^ puffs

To further investigate the effect of diminished IP_3_R1 expression following inhibition of endogenous CREB, we used Total Internal Reflection Fluorescence (TIRF) microscopy to measure the elementary Ca^2+^ signals termed Ca^2+^ puffs, evoked following uncaging of a cell permeable form of caged IP_3_, ci-IP_3_ in hR1 endo cells. These Ca^2+^ signals paradoxically represent a small majority of IP_3_R clusters which are active on stimulation-so called “licensed’’ IP_3_R (Thillaiappan et al., 2017). For this purpose, hR1 endo cells co-transfected with either pcDNA or KCREB and mcherry plasmid were loaded with Cal-520 AM, caged IP_3_ (ci-IP_3_/PM), EGTA-AM and Ca^2+^ puffs were recorded after photolysis of ci-IP_3_ for 40 seconds at a rate of ∼97 frames per second. Ca^2+^ puffs were detected throughout the 40 second period recording following photolysis of ci-IP_3_ in hR1 endo cells transfected with pcDNA or KCREB plasmid (Fig. 7A, B). We specifically imaged only those cells which were transfected with mCherry plasmid (Fig. S4A, B). Interestingly, the number of puff sites per cell were significantly attenuated in hR1 endo cells transfected with KCREB plasmid (65, n=9 cells) when compared to cells transfected with pcDNA (281, n=9 cells) (Fig. 7D) (Videos S1 and S2). Furthermore, similar to the puff sites per cell, the number of puffs per cell remained significantly diminished in KCREB transfected cells (109, n=9 cells) as compared to pcDNA transfected cells (490, n=9 cells) (Fig.7C). The amplitudes (Fig. 7E) and the mean rise (r) and fall (f) times of the Ca^2+^ puffs (Fig. 7F) induced by photolysis of ci-IP_3_ remained similar between hR1 endo cells transfected with pcDNA or KCREB plasmid, indicating that the fundamental biophysical properties of the IP_3_R1 clusters in these cells remained unaltered. These data further substantiates the involvement of CREB in governing the expression of IP_3_R1 which are destined to be active following agonist stimulation.

**Fig. 7:**
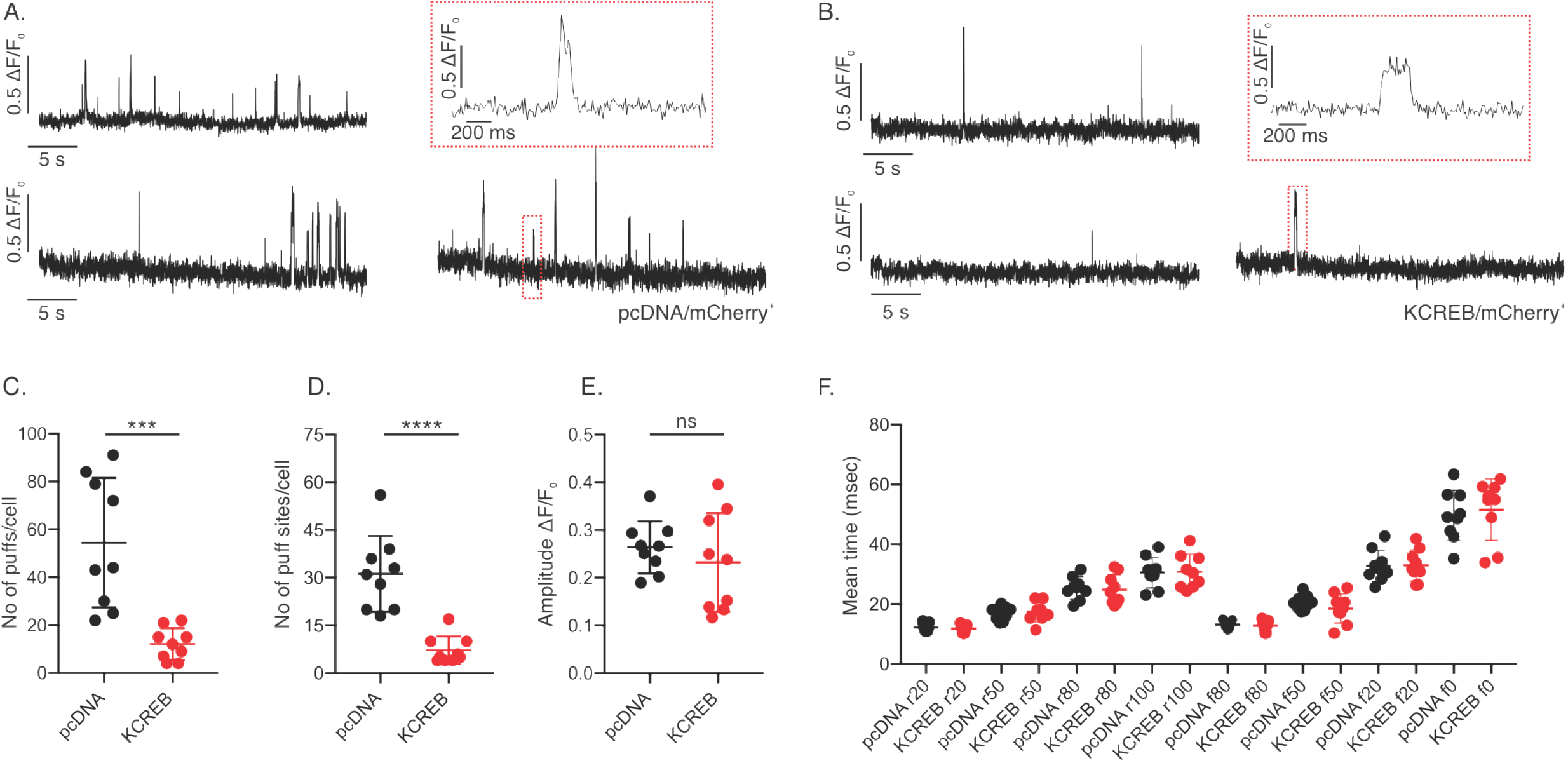
CREB regulates Ca^2+^ puffs produced by IP_3_R1. hR1 endo cells co-transfected with either pcDNA or KCREB and mCherry plasmids (n=9 each) were loaded with Cal-520, ci-IP_3_, EGTA-AM, and imaged using a TIRF microscope. A UV-flash was delivered at 3 seconds to uncage ci-IP_3_. Representative traces of Cal-520 fluorescence ratios (ΔF/F0) from center of single puff sites (1.0 × 1.0 μm) obtained using hR1 endo cells co-transfected with mCherry and either (A) pcDNA or (B) KCREB plasmids. Shown as an inset is a trace (highlighted) on an expanded time scale. (C) The number of puffs were significantly diminished in hR1 endo cells transfected with KCREB plasmid as compared to pcDNA. Similarly, (D) the number of puff-sites were significantly lower in hR1 endo cells transfected with KCREB plasmid compared to cells transfected with pcDNA. (E) The amplitudes of the Ca^2+^ puffs following photolysis of ci-IP_3_ in hR1 endo cells transfected with pcDNA or KCREB plasmid did not differ between these groups. (F) Mean rise (r)- and decay (f)-times for the fluorescence of Ca^2+^ puffs evoked by photolysis of ci-IP_3_ when it increases or decreases to 20%, 50%, 80%, and 100% did not differ between hR1 cells transfected with either pcDNA or KCREB plasmid. Only those cells which were mCherry positive were considered for analyzed. Data presented as mean ± SEM. Statistical significance was determined by Student’s t-test (unpaired, two-tailed). ***p<0.001, ****p<0.0001. ns; not significant.

### KRAP which facilitates proper localization of IP_3_Rs is a target of CREB

Based on the previous reports that KRAP is required for proper localization of IP_3_Rs and in defining their ability to respond to IP_3_ (Fujimoto et al., 2011a), we performed *in silico* analysis to investigate for potential CREB binding sites in the 1 kb proximal promoter region of the KRAP gene. Interesting, our *in silico* analysis using various transcription factor prediction tools revealed putative binding sites for CREB in the proximal promoter domain of KRAP (Table S2). In order to functionally validate the involvement of CREB in regulating the expression of KRAP, we transfected HEK-293 cells with VP16-CREB or KCREB plasmids followed by Western blotting to determine the levels of KRAP. Over-expression of CREB in HEK-293 cells augmented the endogenous KRAP levels (Fig. 8A, B). On the other hand, blocking endogenous CREB using KCREB plasmid diminished the endogenous KRAP level in HEK-293 cells (Fig. 8C, D). These results suggest a potential role for CREB in governing the expression of KRAP and thus contributing to the proper localization and appropriate activity of IP_3_Rs.

**Fig. 8:**
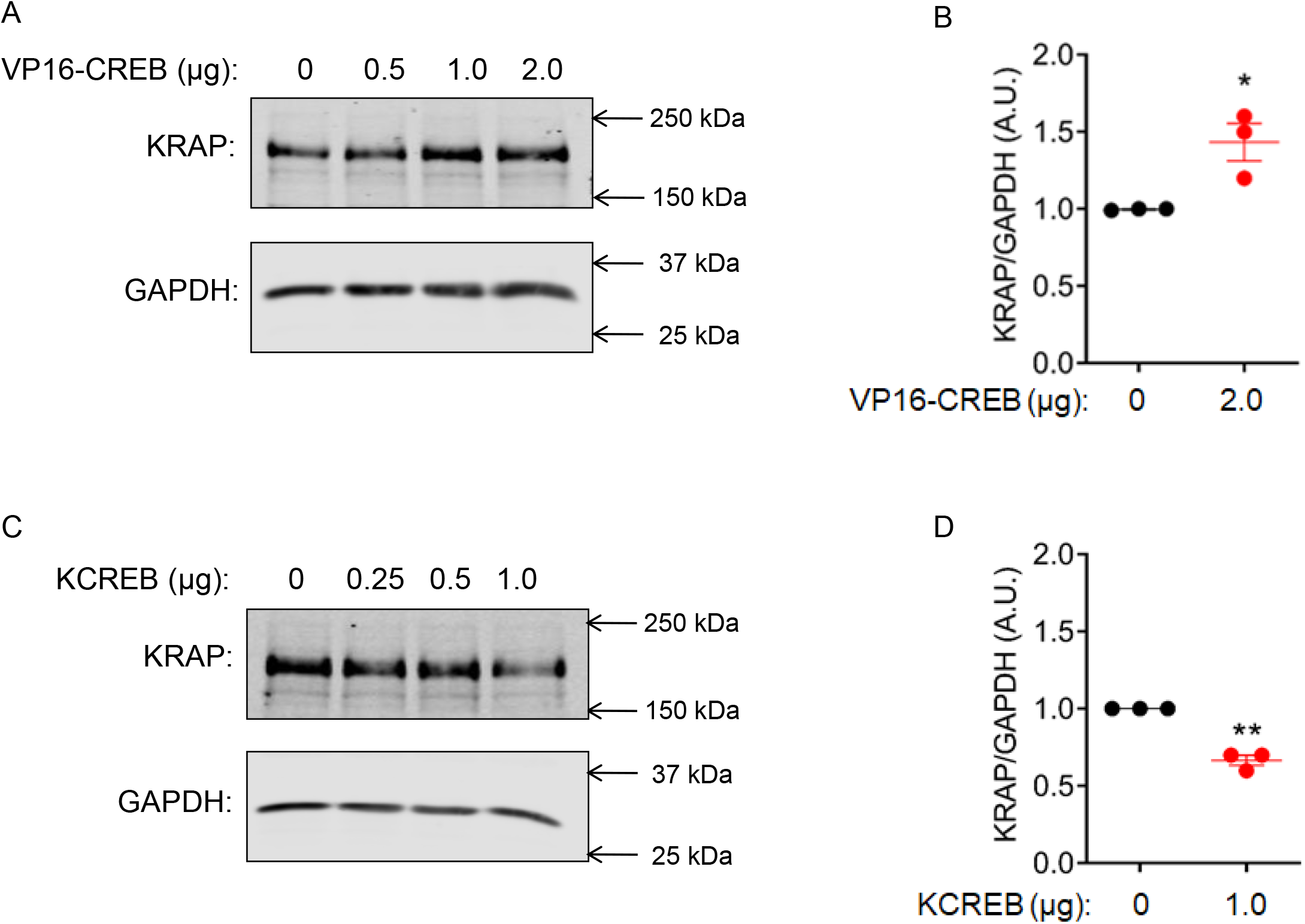
Crucial role of CREB in governing the expression of KRAP. HEK-293 cells were transiently transfected with increasing amounts of VP16-CREB expression plasmid or KCREB plasmid and total protein was isolated 36 hours following transfection. Western blot showing (A) increase in the endogenous KRAP level upon over-expression of CREB in HEK-293 cells. (B) Quantification from three independent blots is shown. (C) Inhibition of CREB, using KCREB, diminished the endogenous KRAP level in HEK-293 cells. (D) Quantification from three independent blots is shown. Statistical significance was determined by Student’s t-test (unpaired, two-tailed). *p<0.05, **p<0.01.

Taken together, our results provide molecular insights into the crucial role of CREB in modulating the expression of IP_3_R1 and KRAP in HEK-293 cells. We speculate that aberrations in the levels of any of these three proteins may be an underlying cause for various pathological conditions.

## DISCUSSION

IP_3_Rs are ubiquitously expressed and often more than one sub-type is present in any given cell type. The functional channel is assembled as a tetramer and can be formed as homo or hetero-tetramers (Foskett et al., 2007). The acute channel activity of the IP_3_R is regulated by a host of regulatory factors including IP_3_, Ca^2+^, ATP, and protein binding partners (Ferris et al., 1990; Foskett and Daniel Mak, 2010; Prole and Taylor, 2016). In addition, IP_3_Rs can be phosphorylated by various kinases. In particular, an early observation was that in response to elevated cAMP levels, IP_3_Rs are phosphorylated by PKA representing a major focus of cross-talk between canonical signaling pathways (Yule et al., 2010). Although the acute effect on channel activity by regulatory factors including PKA phosphorylation has been defined by extensive study, there are relatively few reports of more chronic regulatory events which have the potential to control IP_3_R expression per se. For example, previous reports from our laboratory and other groups have established the roles of Sp1 (Park et al., 2008; Park et al., 2009), AP-1, and NFAT (Mizuno et al., 2012; Mizuno et al., 2015) in regulating IP_3_R1 expression at the transcriptional level. Similarly, the expression of IP_3_R1 is also governed by several cellular factors including TNF-α (Park et al., 2009), and TGF-β (Pacher et al., 2008; Sharma et al., 1997). However, the roles of additional factors influencing the expression of IP_3_R1 remain obscure. As an important consequence of PKA activation in many cell types is to phosphorylate the transcription factor CREB, leading to its nuclear translocation and subsequent expression of target genes. In the current study, we investigated whether IP_3_Rs are targets of CREB transcriptional regulation and whether blocking CREB has any functional consequences on the agonist-induced Ca^2+^ release from IP_3_Rs.

We observed a significant increase in the endogenous IP_3_R1 protein levels upon treatment of HEK-293 cells with 10 µM forskolin for 12 hours. The expression of IP_3_R2 and IP_3_R3 sub-types in these cells, however, remained unaltered (Fig. 1). These observations are supported by evidence from the CREB target gene database (http://natural.salk.edu/CREB/) wherein the binding of CREB to the IP_3_R1 promoter was augmented following treatment with forskolin for 4 hours in HEK-293T cells (binding ratio 1.7) (Zhang et al., 2005). Contrary to our observations in HEK-293 cells, a recent report provided conclusive evidence for involvement of CREB in augmenting the expression of IP_3_R2 levels upon fasting or treatment of hepatocytes with forskolin (Kruglov et al., 2017). Such a disparity may be attributed to differences in epigenetic modifications in these cell types. We conclude that IP_3_Rs may be differentially regulated by CREB in a cell type-or tissue-specific manner.

Next, our *in silico* predictions using various transcription factor binding prediction tools revealed putative binding sites for CREB in the 1 kb proximal promoter regions of all the three IP_3_R sub-types. Moreover, RNAseq data mined from GTEx portal revealed a significant positive correlation between the transcript levels of IP_3_R1/2/3 and CREB across various human tissues (n=54) underscoring a potential role for CREB in regulating the expression of all the three IP_3_R sub-types (Fig. 2). However, over-expression of CREB in HEK-293 cells resulted in a dose-dependent increase in the endogenous IP_3_R1 protein levels without any effect on the expression of IP_3_R2 and IP_3_R3 sub-types. Conversely, blocking CREB caused a decrease in IP_3_R1 levels in HEK-293 cells (Fig. 3). Interestingly, the IP_3_R1 promoter harbors a TATA box while both the IP_3_R2 and IP_3_R3 promoters lack a typical TATA box (Konishi et al., 1997). Initial screening showed that CREB binds to the proximal promoter regions containing a TATA box (Zhang et al., 2005); however, later investigations provide evidence that CREB could bind to distal promoter regions as well (Everett et al., 2013). This is further supported by the fact that IP_3_R2 (which lacks a typical TATA box) is still regulated by CREB (Kruglov et al., 2017). Moreover, it is well known that CREB is not recruited on methylated promoters (Zhang et al., 2005) indicating that the IP_3_R2 and IP_3_R3 promoters may be methylated in HEK-293 cells.

CREB is a ubiquitously expressed transcription factor. During fasting, glucagon via PKA promotes Ser-133 phosphorylation of CREB. CREB in turn augments the expression of gluconeogenic gene including pyruvate carboxylase (PC), phosphoenolpyruvate carboxykinase 1 (PEPCK1) and glucose-6-phosphatase (G6PC), Peroxisome proliferator-activated receptor-gamma coactivator (PGC-1alpha) (Koo et al., 2005). Of note, glucagon via PKA also phosphorylates IP_3_R1 which results in increased cytosolic Ca^2+^. This increase in cytosolic calcium promotes CRTC2 dephosphorylation and nuclear localization which in turn enhances the expression of gluconeogenic genes (Wang et al., 2012). Furthermore, glucagon, in a transcriptional-independent manner via IP_3_R1 promotes gluconeogenesis (Perry et al., 2020). Strikingly, a transgenic mouse line (D2D) characterized by disruption in the *Itpr1* loci and the *Itpr1* heterozygous mutant mice (opistotonus mouse), opt/+, exhibit glucose intolerance supporting a role for IP_3_R1 in glucose homeostasis (Ye et al., 2011). We observed a decrease in IP_3_R1 protein levels following inhibition of PKA in HEK-293 cells and blunted expression of IP_3_R1 in response to forskolin upon blocking CREB (Figs. 3 and 4). Our results indicate that PKA in addition to its role in phosphorylating IP_3_R1 also appears to govern the expression of IP_3_R1. All these evidence strongly indicate IP_3_R1 may act in a positive feed-back manner via CREB to regulate their own expression.

Does blocking the function of CREB have any effect on agonist-induced Ca^2+^ release from IP_3_R1 receptors? Indeed, transient transfection of KCREB plasmid in a population-based assay revealed a significant decrease in agonist-induced Ca^2+^ release from hR1 endo cells. Such a decrease in agonist-induced Ca^2+^ release was not observed in KCREB transfected hR3 endo cells (Fig. 5). We verified these results by performing single cell imaging using hR1 endo cells transfected with the KCREB plasmid (Fig. 6). These observations confirm the functional implications of CREB on agonist-induced Ca^2+^ release mediated by IP_3_R1 sub-type.

Research in the past decade revealed that all three IP_3_Rs, to a similar extent, can evoke elementary Ca^2+^ signals called “Ca^2+^ puffs” (Lock et al., 2018; Mataragka and Taylor, 2018). Each cell type expresses numerous IP_3_Rs most of which are mobile and diffuse freely in the endoplasmic reticulum. In response to various stimuli or upon uncaging ci-IP3, Ca^2+^ puffs arise repeatedly from a small fraction of localized and immobile IP_3_Rs. These clusters of IP_3_Rs are termed as ‘licensed’ IP_3_Rs (Thillaiappan et al., 2017). We investigated the effect of blocking CREB in hR1 endo cells on the Ca^2+^ puffs using TIRF microscopy. Interestingly, we observed a significant decrease in the number of puffs and puff sites following uncaging ci-IP3/PM upon blocking CREB in hR1 endo cells compared to control cells (Fig. 7). These results underscore the functional consequences of blocking CREB in hR1 endo cells.

KRAP physically associates with the amino-terminal residues of IP_3_Rs and co-immunoprecipitates with IP_3_Rs both *in vivo* and also in various cell lines including HEK-293 cells (Fujimoto et al., 2011a). KRAP may play an important role in ensuring proper immobilization and licensing of IP_3_Rs. Furthermore, siRNA mediated knockdown of KRAP diminished IP_3_R-mediated calcium release (Fujimoto et al., 2011b). We observed an increase or decrease in KRAP levels upon over-expression or inhibition of CREB (Fig. 8). These findings suggest that in addition to regulating the expression of IP_3_R1, CREB, by modulating the expression of KRAP, plays a vital role in proper localization and licensing of IP_3_R1 in HEK-293 cells.

In conclusion, this preliminary study using HEK-293 cells as a model provides evidence indicating a crucial role for CREB in governing not only the expression, but proper localization, and thus licensing of IP_3_R1 which are destined to respond to various cues (Fig.9). We speculate that dysregulation of CREB-mediated IP_3_R1 regulation may have implications in numerous pathophysiological conditions which remain to be further investigated.

**Fig. 9:**
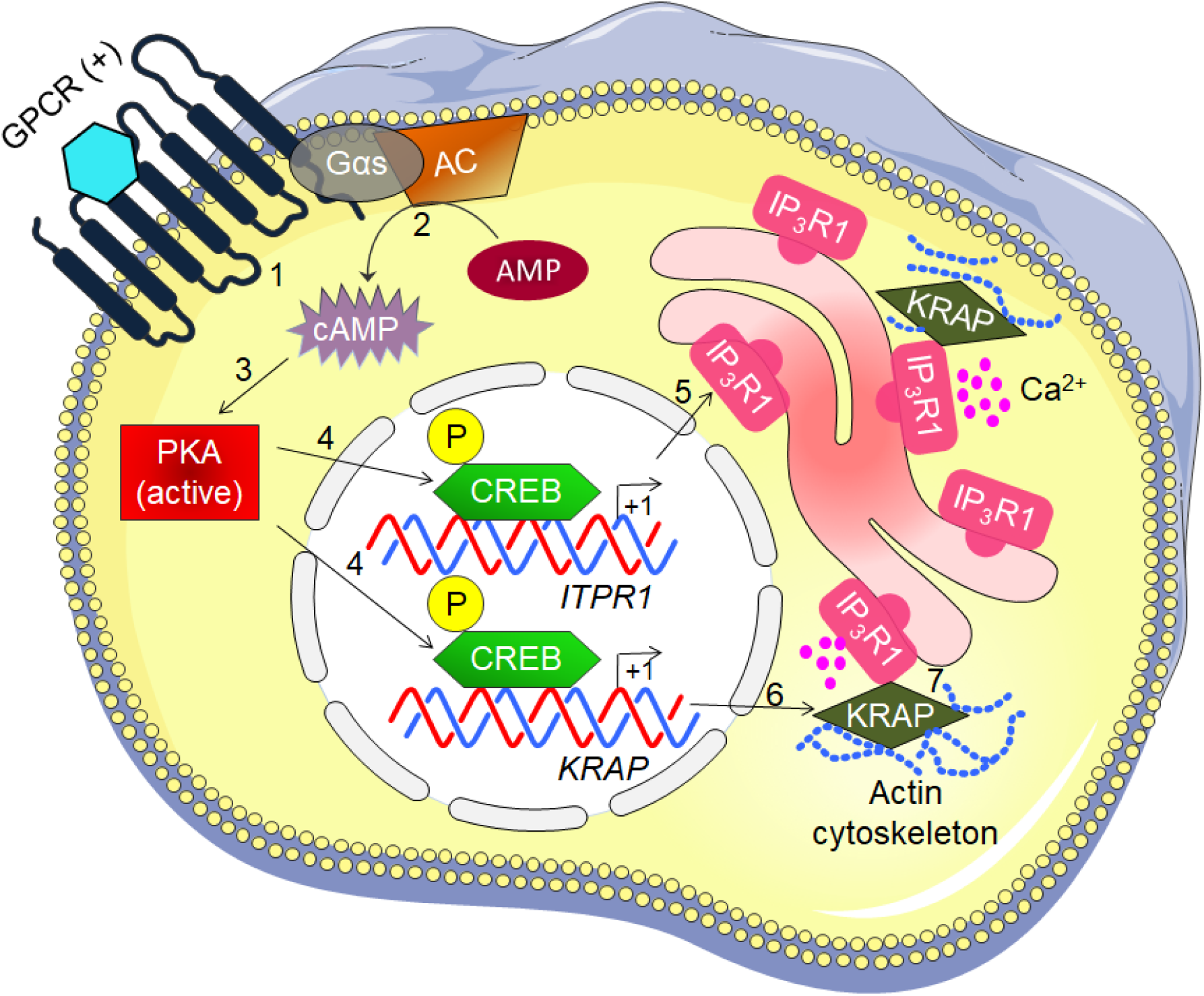
A summary depicting the involvement of PKA-CREB axis in augmenting the expression of IP_3_R1 in HEK-293 cells. Activated GPCR (1) augment the intracellular levels of cAMP (2). cAMP promotes activation of PKA (3). PKA in turn phosphorylates CREB (4) which enhances the endogenous protein levels of IP_3_R1 (5) and KRAP (6). KRAP physically associates with IP_3_R1 (7) ensuring proper localization and licensing of IP_3_R1. GPCR, G protein-coupled receptors; Gαs, Gs alpha subunit; AC, adenylate cyclase; AMP, adenosine monophosphate; cAMP, cyclic AMP; PKA, protein kinase A; CREB, cAMP response element-binding protein; P, phosphorylation; IP_3_R1, inositol 1,4,5 trisphosphate receptor type 1; KRAP, KRAS-Induced Actin-Interacting Protein; (+), activate; Ca^2+^, calcium.

## MATERIALS AND METHODS

### Plasmids

VP16-CREB expression plasmid, a constitutively active form of CREB, was kindly provided by Dr. David Ginty (Harvard Medical School, USA). KCREB plasmid, a dominant-negative form of CREB, was kindly provided by Dr. Richard Goodman (Vollum Institute, USA). pRSV-PKI-v2 - PKI plasmid was obtained from Addgene (Catalog #45066). pcDNA 3.1+ was procured from Invitrogen, USA. mCherry-3HA plasmid was generated in our laboratory.

### *In silico* predictions and GTEx data

The sequences for upstream 1 kb proximal promoter regions of ITPR1 (NM_001099952), ITPR2 (NM_002223), ITPR3 (NM_002224), and KRAP (NM_006751) were retrieved from UCSC genome browser (https://genome.ucsc.edu/). Predictions for putative transcription factor binding sites in the proximal promoter domains were performed using Alibaba2.1, TFBIND, ConSite, and PROMO tools. For correlation analysis between the transcript levels of ITPRs and CREB across various tissues, RNAseq data was mined from GTEx portal (https://gtexportal.org/home/).

### Cell culture and transient transfections

HEK-293, hR1 endo, and hR3 endo cells were cultured in Dulbecco’s modified Eagle’s medium (DMEM) supplemented with 10% fetal bovine serum, 100 U/ml penicillin, and 100 μg/ml streptomycin in 10 cm^2^ cell culture dish. For transient transfections, cells were seeded in appropriate cell culture dishes. Following 24 hours seeding, the cells were transfected with indicated amounts of pcDNA, VP16-CREB, KCREB, or PKI plasmids using Lipofectamine 2000 Transfection Reagent (Thermo Fisher Scientific, USA) according to the manufacturer’s instructions. pcDNA was used as a balancing plasmid to transfect equal amount of DNA in each well. Similarly, the hR1 or hR3 endo cells were transfected with pcDNA or KCREB prior to imaging. In certain instances, the cells were co-transfected with mCherry plasmid at a 1:10 ratio (mCherry:plasmid of interest, respectively). The cells were left in an incubator for 36 hours post transfection before proceeding for Western blotting or Ca^2+^ imaging.

### Western Blotting

For Western blotting, total protein was isolated from HEK-293 control cells or cells treated with 10 μM forskolin for 12 hours using membrane-bound extraction buffer (10 mM Tris-HCl, 10 mM NaCl, 1 mM EGTA, 1 mM EDTA, 1 mM NaF, 20 mM Na_4_P_2_O_7_, 2 mM Na_3_VO_4_, 1% Triton X-100 (v/v), 0.5% sodium deoxycholate (w/v), and 10% glycerol) supplemented with protease inhibitors. Similarly, total protein was also isolated from cells transfected with pcDNA, CREB, KCREB, or PKI plasmids 36 hours post-transfection. In order to determine the involvement of CREB in mediating the effects of forskolin, cells were treated with forskolin 24 hours post-transfection with pcDNA or KCREB and left for an additional 12 hours before protein isolation. Briefly for protein isolation, following addition of appropriate amount of lysis buffer to cells, the cells were harvested in 1.5 mL tubes, and placed on ice for 30 minutes. In order to disrupt the pellet, the tubes were vortexed for 10 seconds every 10 minutes and returned on ice. Following 30 minutes incubation on ice, the cells lysates were centrifuged at 13,000 rpm at 4°C for 10 minutes. The supernatant was transferred to new labeled tubes. Protein concentration in the lysates was estimated using Dc protein assay kit (Bio-Rad). The lysates (30 μg) were then subjected to SDS-PAGE and transferred to a nitrocellulose membrane. The membranes were incubated with indicated primary antibodies and appropriate secondary antibodies before imaging with an Odyssey infrared imaging system (LICOR Biosciences). Band intensities were quantified using Image Studio Lite Ver 5.2 and presented as ratios of IP_3_Rs/KRAP to GAPDH.

### Population based Ca^2+^ imaging

hR1 or hR3 endo cells in 10 cm^2^ cell culture dish were transfected with 5 μg pcDNA or KCREB plasmid. Following 36 hours transfection, the cells were loaded with 4 μM Fura-2/AM in cell culture media and left in dark for an hour. The cells were subsequently washed with imaging buffer (10 mM HEPES, 1.26 mM Ca^2+^, 137 mM NaCl, 4.7 mM KCl, 5.5 mM glucose, 1 mM Na_2_HPO_4_, 0.56 mM MgCl_2_, at pH 7.4). Equal number (300,000 cells/ well) of pcDNA and KCREB transfected cells were seeded into each well of a black-walled 96-well plate. The cells were centrifuged at 200g for 2 minutes to plate the bottom of the wells and left in an incubator for another 30 minutes before imaging. Fura-2/AM imaging was carried out by alternatively exciting the loaded cells between 340 nm and 380 nm; emission was monitored at 510 nm using FlexStation 3 (Molecular Devices). Peak response to various concentrations of CCh (0.1-100 μM) was determined using SoftMax© Pro Microplate Data Acquisition and Analysis software as described previously (Terry et al., 2020). Data from at least three individual plates after curve fitting using a logistic dose-response equation in GraphPad Prism 8 are presented.

### Single cell Ca^2+^ imaging

hR1 endo cells seeded on 15 mm glass coverslips in 35 mm dishes were co-transfected with pcDNA or KCREB and mCherry plasmids. Following 36 hours transfection, the cells were washed with imaging buffer. The glass coverslip was attached to a Warner perfusion chamber using vacuum grease. The cells were incubated with 2 μM Fura-2/AM for 25 minutes in the dark at room temperature for loading. Cell were then perfused with imaging buffer and stimulated with indicated concentrations of CCh. Ca^2+^ imaging was performed using an inverted epifluorescence Nikon microscope equipped with a 40 x oil immersion objective. Fura-2/AM imaging was carried out by alternatively exciting the loaded cells between 340 nm and 380 nm; emission was monitored at 505 nm. Images were captured every second with an exposure of 20 ms and 4 × 4 binning using a digital camera driven by TILL Photonics software as previously described (Terry et al., 2020). Image acquisition was performed using TILLvisION software and data was exported to Microsoft excel. Experiments were repeated at least three times and only cells positive for mCherry were analyzed.

### Detection and analysis of Ca^2+^ puffs using TIRF microscopy

hR1 endo cells were grown on 15-mm glass coverslips coated with poly-D-lysine (100 μg/ml) in a 35-mm dish for 24 hours were co-transfected with either 1 μg pcDNA or KCREB and 100 ng mCherry plasmids. Following 36 hours transfection, prior to imaging, the cells were washed three times with imaging buffer. The cells were subsequently incubated with Cal520-AM (5 µM; AAT Bioquest #21130) and ci-IP_3_/PM (1 μM, Tocris #6210) in imaging buffer with 0.01 % BSA in dark at room temperature. After 1 hour incubation, the cells were washed three times with imaging buffer and incubated in imaging buffer containing EGTA-AM (5 μM, Invitrogen #E1219). After 45 minutes incubation, the media was replaced with fresh imaging buffer and incubated for additional 30 minutes at room temperature to allow for de-esterification of loaded reagents.

Following loading, the coverslip was mounted in a chamber and imaged using an Olympus IX81 inverted total internal reflection fluorescence microscopy (TIRFM) equipped with oil-immersion PLAPO OTIRFM 60x objective lens/1.45 numerical aperture. Olympus CellSens Dimensions 2.3 (Build 189987) software was used for imaging. mCherry positive cells were identified using a 561 nm laser. Subsequently, the cells were illuminated using a 488 nm laser to excite Cal-520 and the emitted fluorescence was collected through a band-pass filter by a Hamamatsu ORCA-Fusion CMOS camera. The angle of the excitation beam was adjusted to achieve TIRF with a penetration depth of ∼140 nm. Images were captured from a final field of 86.7 µm x 86.7 µm (400 x 400 pixels, one pixel=216 nm) at a rate of ∼97 frames/second (binning 2 x 2) by directly streaming into RAM. To photorelease IP_3_, UV light from a laser was introduced to uniformly illuminate the field of view. Both the intensity of the UV flash and the duration (1 second) for uncaging IP_3_ were optimized to prevent spontaneous puff activity in the absence of loaded ci-IP_3_. Images were exported as vsi files. Images, 5 seconds before and 40 seconds after flash photolysis of ci-IP_3_, were captured, as described previously (Emrich et al., 2021).

The vsi files were converted to tiff files using Fiji and further processed using FLIKA, a Python programming-based tool for image processing (Ellefsen et al., 2014). From each recording, ∼300 frames (∼3 seconds) before photolysis of ci-IP_3_ were averaged to obtain a ratio image stack (F/Fo) and standard deviation for each pixel for recording up to 30 seconds following photolysis. The image stack was Gaussian-filtered, and pixels that exceeded a critical value (1.0 for our analysis) were located. The ‘Detect-puffs’ plug-in was utilized to detect the number of clusters (puff sites), number of events (number of puffs), amplitudes and durations of localized Ca^2+^ signals from individual cells. All the puffs identified automatically by the algorithm were manually confirmed before analysis. The results from FLIKA were saved as excel and graphs were plotted using GraphPad Prism 8.

### Statistical Analysis

All statistical tests were conducted using GraphPad Prism 8 and data is presented as mean ± SEM. Statistical significance was determined by student’s t-test (unpaired, two-tailed) or by one-way ANOVA with Tukey’s multiple comparisons post-test. *p<0.05, **p<0.01, ***p<0.001,****p<0.0001 with respect to control condition. ns; not significant.

## Acknowledgements

The authors wish to thank Dr. David Ginty (Harvard Medical School, USA) for providing the VP16-CREB expression plasmid construct. The authors also wish to thank Dr. Richard Goodman (Vollum Institute, USA) for providing the KCREB plasmid construct. The authors thank all the members of the lab especially Dr. Sundeep Malik and Ms. Kai-Ting Huang for their valuable suggestions.

## Competing interests

The authors of this work have no competing interests to disclose.

## Funding

This work was supported by National Institutes of Health Grant R01 DE014756 to Dr. David I. Yule.

## Notes

### Competing Interest Statement

The authors have declared no competing interest.

